# Temporal dynamics of functional networks in long-term infant scalp EEG

**DOI:** 10.1101/2020.09.21.307082

**Authors:** Rachel J. Smith, Ehsan Alipourjeddi, Cristal Garner, Amy L. Maser, Daniel W. Shrey, Beth A. Lopour

## Abstract

Human functional connectivity networks are modulated on time scales ranging from milliseconds to days. Rapid changes in connectivity over short time scales are a feature of healthy cognitive function, and variability over long time scales can impact the likelihood of seizure occurrence. However, relatively little is known about modulation of healthy functional networks over long time scales. To address this, we analyzed functional connectivity networks calculated from long-term EEG recordings from 19 healthy infants. Networks were subject-specific, as inter-subject correlations between weighted adjacency matrices were low. However, within individual subjects, both sleep and wake networks were stable over time, with stronger functional connectivity during sleep than wakefulness. This enabled automatic separation of wakefulness and sleep states via principle components analysis of the functional network time series, with median classification accuracy of 91%. Lastly, we found that network strength, degree, clustering coefficient, and path length significantly varied with time of day, when measured in both wakefulness and sleep. Together, these results suggest that modulation of healthy functional networks occurs over long timescales and is robust and repeatable. Accounting for such temporal periodicities may improve the physiological interpretation and use of functional connectivity analysis to investigate brain function in health and disease.

## 1. Introduction

Resting-state functional connectivity can be assessed with a variety of imaging modalities, including scalp electroencephalography (EEG) (Burroughs, Morse, Mott, & Holmes, 2014; Shrey et al., 2018), intracranial EEG (M. A. Kramer et al., 2011), magnetoencephalography (MEG) (Stam, Nolte, & Daffertshofer, 2007), and functional magnetic resonance imaging (fMRI) (Allen, Damaraju, Eichele, Wu, & Calhoun, 2018; Chen, Feng, Zhao, Yin, & Wang, 2008; Damoiseaux et al., 2006; De Asis-Cruz, Bouyssi-Kobar, Evangelou, Vezina, & Limperopoulos, 2015; Fingelkurts & Fingelkurts, 2011). In fMRI, the brain’s metabolic activity, as measured with the blood oxygen level-dependent (BOLD) signal, describes network states that are activated during rest and differentiates persistent states from task-dependent activity (Damoiseaux et al., 2006; De Asis-Cruz et al., 2015). Although it is the most popular modality to investigate resting-state functional connectivity in healthy subjects, the low temporal resolution of fMRI limits the comparison to networks derived from other modalities, unless dual acquisition of the neural signals is possible (Chang, Liu, Chen, Liu, & Duyn, 2013; Sadaghiani et al., 2010). Additionally, most studies of fMRI-based functional connectivity are assessed with 10-20 minutes of data, and studies lasting hours or days are infeasible. This precludes analysis of the long-term dynamical changes in functional connectivity.

EEG-based functional connectivity provides an alternative to fMRI that enables long-term recordings of spontaneous brain activity (M. A. Kramer et al., 2011). Long-term, ultradian periodicities in EEG signal features, such as frequency band power, have been well studied (Aeschbach et al., 1999; Kaiser, 2008). For example, ultradian modulation of EEG power has been shown to occur in the 9-11 Hz frequency band over 2 hour periods, in the 11-13 Hz frequency band over 4 hour periods, and in the fronto-centrally located upper beta band over 3-4 hour periods (Kaiser, 2008). Such periodicities have not been as extensively studied in functional networks. EEG-based functional networks are paradoxically stable on the time scale of minutes, despite the visual variability in EEG time series (Chapeton, Inati, & Zaghloul, 2017; M. A. Kramer et al., 2011), and the networks are unique to individual subjects (Chu et al., 2012; Shrey et al., 2018). Specifically, changes in the small-worldedness of the functional networks were found during the transitions from wakefulness to sleep (Ferri, Rundo, Bruni, Terzano, & Stam, 2008). In intracranial EEG data, functional network modulation over the course of days exhibited strong circadian effects in graph theoretical measures of the networks, such as the clustering coefficient and path length (Geier, Lehnertz, & Bialonski, 2015; Kuhnert, Elger, & Lehnertz, 2010). However, the impact of circadian modulation on the structure, stability, and robustness of scalp EEG-based functional networks remains unknown.

In addition to modulating healthy cognitive function, the circadian rhythm can affect brain dysfunction. For example, long-term functional network changes have been used to predict seizure onset (Anastasiadou et al., 2016; Baud et al., 2018; Karoly et al., 2017; Kuhnert et al., 2010), and functional network properties were better predictors of seizure onset than signal power or other signal properties (Mitsis et al., 2017). Additionally, the physiological changes in the functional networks associated with the time of day have a far greater impact than changes in the networks due to seizure onset, highlighting the need for studies to account for these physiological changes when creating seizure prediction models (Kuhnert et al., 2010; Mitsis et al., 2017; Schelter, Feldwisch-Drentrup, Ihle, Schulze-Bonhage, & Timmer, 2011).

Therefore, to quantify normal physiological changes in brain networks over long time scales, we measured EEG-based functional connectivity in long-term recordings from healthy infants. Infant EEG provides a unique advantage over adult recordings, as infants sleep more frequently during the day, allowing us to capture more sleep/wake state transitions than would have been possible in an adult study. This also enabled us to disentangle the changes in the functional connectivity network that were due to sleep/wake transitions, as opposed to circadian changes that were associated with a particular time of day. Here, we describe the functional networks associated with sleep and awake states, assess the inter-subject variability in those state-specific networks, and identify graph theoretical measures that separate the two states. Then we show that measurements of the sleep/wake networks in individual subjects are stable and repeatable, enabling robust classification of the states over many hours of data using principle components analysis. Lastly, we show circadian variation in functional connectivity strength and graph theoretical measures. This work increases our understanding of the brain’s physiological fluctuations in functional connectivity, which has the potential to inform investigations of functional networks that are disrupted due to pathology.

## 2. Methods

### 2.1 Subject recruitment and EEG recording

This prospective study was approved by the Institutional Review Board of the Children’s Hospital of Orange County (CHOC). Subjects were recruited and consented from June 2017 to February 2019 and underwent overnight long-term video EEG recording to rule out a form of pediatric epilepsy called infantile spasms. If the infant was not diagnosed with infantile spasms, they were classified as a control subject. Clinical data were collected at the time of enrollment. Subjects were deemed “healthy” controls if they: (1) Exhibited a normal EEG recording, (2) did not receive a diagnosis of epilepsy, (3) had no known neurological conditions, and (4) were developmentally normal for age (as assessed with the Vineland Adaptive Behavior Scales, 3^rd^ Edition (Sparrow, Cicchetti, & Saulnier, 2016)). EEG data were sampled at 200 Hz with impedances below 5 kΩ.

A certified sleep technologist at the Children’s Hospital of Orange County manually delineated time periods of wakefulness, rapid eye movement (REM) sleep, and non-REM sleep stages (N1, N2, N3) in all EEG recordings in accordance with the American Academy of Sleep Medicine (AASM) guidelines. For our analysis, time periods of sleep and wakefulness were separated based on these markings. For comparison with automatic sleep staging (see Section 2.6), we combined N1, N2, N3, and REM sleep stages into one “sleep” category.

### 2.2 EEG Pre-processing

EEG data were re-referenced offline to the common average. Artifactual time periods were identified with an automatic extreme value artifact detector, similar to previously published methods (Durka, Klekowicz, Blinowska, Szelenberger, & Niemcewicz, 2003; Moretti et al., 2003). Specifically, to identify artifacts we broadband bandpass filtered the data (1.5-40 Hz, Butterworth filter, chosen to match the settings of clinical EEG viewing/analysis), subtracted the mean from each channel, and calculated the standard deviation of each zero-mean time series. Artifacts were defined as time points in which the absolute value of the voltage exceeded a threshold of 7.5 standard deviations above the mean value in any single channel. We chose this threshold because it resulted in the best correspondence between automatically-detected and visually-identified artifacts in a previous dataset (R.J. Smith et al., 2017). To ensure that the entire artifact was marked, a buffer of 0.9 seconds was added to both sides of each contiguous set of time points containing extreme amplitude values. Data recorded during EEG impedance checks were also marked as artifact. For the connectivity analysis, a broadband bandpass-filter was applied to the raw, re-referenced data (0.5-55Hz, Butterworth filter). One-second epochs that contained artifactual data were removed from all channels after filtering.

### 2.3 Functional connectivity

We calculated functional connectivity networks via cross-correlation using the method developed by Kramer et al. (Mark A. Kramer, Eden, Cash, & Kolaczyk, 2009) and Chu et al. (Chu et al., 2012) and previously applied to infant EEG data in (Shrey et al., 2018). We chose cross-correlation because it is a simple bivariate measure that is highly sensitive to linear changes in EEG activity (David, Cosmelli, & Friston, 2004; Jalili, Barzegaran, & Knyazeva, 2014) and has been shown to be comparable to other measures of synchronization (Jalili et al., 2014; Quian Quiroga, Kraskov, Kreuz, & Grassberger, 2002). Although cross-correlation is generally insensitive to nonlinear interactions in the EEG, we opted for this rapid and straightforward linear measure of synchronization because no nonlinear metric has been shown to reliably measure actual changes in coupling strength while discounting spurious increases in due to changes in other signal properties (David et al., 2004; Pereda, Rial, Gamundi, & González, 2001).

The functional connectivity calculation was performed as described in (Shrey et al., 2018); we briefly summarize it here. Data were divided into one-second epochs, and the EEG signals in each epoch (one from each channel) were normalized to have zero-mean and unit variance. For each epoch, we calculated the cross-correlation between every pair of channels and identified the maximum of the absolute value of the cross-correlation. Epochs in which the maximal cross-correlation value occurred at zero time lag were excluded, as they were likely a result of volume conduction (Chu et al., 2012). A partial correlation with the common-average reference time series was performed to test whether the reference induced the correlation measured between the channels (Shrey et al., 2018). If there was a large difference between the partial correlation and the correlation value between the channels, the measured correlation was presumed to have resulted from referencing artifact and the epoch was removed from further analysis. Z-values were calculated for the non-artifactual epochs by dividing the Fisher-transformed correlation coefficient value by the estimated standard deviation, taking the autocorrelation of each channel epoch into account (Chu et al., 2012; Mark A. Kramer et al., 2009). The z-values were compared to a baseline distribution created via permutation resampling. Permutation resampling was performed by selecting two random one-second epochs of data from the time series that were separated by at least one second, calculating the correlation between the channels, and iterating this procedure 500 times (Nichols & Holmes, 2003). The standardized correlation values from all iterations were sorted and the threshold of significance was defined as the value corresponding to the 95^th^ percentile of the distribution for each electrode pair. For each epoch, correlation values between channel pairs that exceeded this threshold value were deemed to be significant, and these connections were assigned a value of one; connections that did not exceed this threshold were assigned a value of zero. Thus for *p* EEG channels, the output of the connectivity calculation for each epoch was a binary matrix of dimension *p* × *p*.

Across epochs, connectivity data were stored in a three-dimensional array ***Q**,* where the binary value at position ***Q***(*i,j, k*) represented the connection between electrode *i* and electrode *j* in epoch *k*. Then the overall connection strength between two channels was calculated as the fraction of time series epochs in which there was a significant connection between them. For wakefulness, this calculation used the *N_w_* binary connectivity matrices that coincided with times of wakefulness based on manual sleep staging to obtain ***Q***_*w*_(*i,j*) = (1/*N_W_*) ∑_*k*∈_wake_*Q*(*i,j,k*)_. We also performed the analogous calculation for ***Q**_s_* using the *N_s_* epochs marked as sleep (N1, N2, N3, and REM). Thus, we calculated one average wakefulness network ***Q**_w_* and one average sleep network ***Q**_s_* for each subject. We evaluated network strength by applying a proportional threshold to the connectivity values (Garrison, Scheinost, Finn, Shen, & Constable, 2015). Specifically, we calculated the average of the strongest 10% of connections in the wakefulness and sleep networks. The Benjamini-Hochberg procedure was used to correct for multiple comparisons where applicable (Benjamini & Hochberg, 1995).

### 2.4 Graph theoretical measures

Studies of functional connectivity are often augmented by complex network analysis because it provides easily computable measures with biophysiological significance (Rubinov & Sporns, 2010). We computed three weighted graph theoretical measures that summarize the functional connectivity networks computed in this study. First, we calculated the *degree* for each node by summing the weights of the connections incident to that node (Bullmore & Sporns, 2009; Rubinov & Sporns, 2010). Second, the *clustering coefficient* is defined as the fraction of a node’s neighbors that are also neighbors of each other and is thought to quantify the level of functional segregation in the brain network (Rubinov & Sporns, 2010; Watts & Strogatz, 1998). In the weighted network case, the clustering coefficient is derived from the “intensity” and “coherence” of a subgraph using measures of its geometric and arithmetic mean (Onnela, Saramäki, Kertész, & Kaski, 2005). Before calculating the clustering coefficient, we normalized the adjacency matrix by dividing each element by the maximum connection value in the network (Antoniou & Tsompa, 2008; Onnela et al., 2005). Third, we computed the *shortest path length,* which reports the minimum sum of the edge “lengths” for a path from one node to another (Antoniou & Tsompa, 2008; Rubinov & Sporns, 2010; Stam & Reijneveld, 2007). In our case, because we assume that information flow will increase with higher connection values, we defined the edge lengths as the inverse of the edge weights. Thus, the minimum sum of these inverse edge weights maximizes the connectivity strength between each pair of electrodes (Rubinov & Sporns, 2010). The shortest path length is one of the most common metrics to assess functional integration (Rubinov & Sporns, 2010). Similar to the clustering coefficient calculation, we normalized the adjacency matrix before computing the inverse and determining the shortest paths between nodes in the network (Antoniou & Tsompa, 2008; Onnela et al., 2005).

### 2.5 Time-varying functional connectivity measurement

To analyze time-varying changes in the functional connections, we averaged the binary *p* × *p* matrices across windows of successive epochs. Let ***Q**_n_* represent the *p* × *p* matrix averaged over a window of *n* one-second epochs. The value of ***Q_n_***(*i,j*) indicates the proportion of epochs in which the connection between channel *i* and channel *j* was significant, analogous to the values of ***Q**_w_* and ***Q**_s_* for wakefulness and sleep, respectively. For our analysis, we calculated ***Q*_300_**, the averaged connectivity matrix in a window of 300 seconds, and this window was shifted in 30-second increments (90% overlap). We chose a window size of 300 seconds because networks were shown to be stable over this amount of time in two separate studies (Chu et al., 2012; Shrey et al., 2018).

### 2.6 Classification of network states

We hypothesized that different brain states, e.g., sleep or wakefulness, would be associated with different functional networks. However, we also expected that the functional network associated with a single state would be consistent over time. We used principle components analysis (PCA) to obtain the latent variable that best distinguished wakefulness and sleep, as we assumed this state transition would be the greatest source of variance in the functional connectivity network.

To perform PCA on the functional connectivity networks over time, we first calculated ***Q*_300_** in 300-second windows with 90% overlap, as described in Section 2.5. We then placed the values for all unique connections in ***Q*_300_** (171 channel pairs in total, excluding self-connections) into a column vector *c* and normalized it to have zero mean and unit variance. We concatenated *m* successive vectors, where *m* denotes the number of windows that were available in the dataset, into matrix *C* and subtracted the mean from each row to ensure that the distribution of connections for each channel pair was zero-mean. We performed PCA on the normalized functional connectivity time series *C* to ascertain the latent networks that explained the most variance in the data. We then calculated the time course of the first principle component, which represents the relative weight assigned to that component as a function of time.

Because we hypothesized that this time series would exhibit transitions between two different states, likely representing wakefulness and sleep, we fit a two-component gaussian mixture model (GMM) to the principle component time series. We then calculated normalized probability distribution functions (PDFs) for the two GMM distributions. The threshold to separate the states was defined as the intersection of the two PDFs. To avoid finding intersections at the extreme tails of the distributions, we calculated the PDF ratio and identified the index where this ratio was closest to 1:

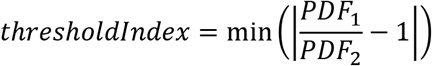

The principle component value associated with this index was the threshold that best distinguished the two states. We therefore used this value to classify the networks from all time points into two states, and we compared these results to ground truth visual sleep scoring done by a certified EEG sleep technician.

### 2.7 Calculation of network stability

We assessed stability of the functional connectivity measurement by performing 2-dimensional correlations between independent average connectivity networks during wakefulness or sleep. We first concatenated all epochs during sleep (*N_s_*) or wakefulness (*N_w_*) and then calculated *N_s_/n* or *N_w_/n* sequential, nonoverlapping measurements of ***Q_n_*** where *n* is the size of the window. Then a 2-D correlation was calculated between each successive measurement of ***Q_n_*** with the MATLAB function “corr2(),” and this was repeated for window sizes ranging from 10 seconds to 200 seconds. The mean of the correlation values was recorded for each window size for each subject. The mean and 95% confidence interval of the average correlation coefficient values for all subjects were then plotted as a function of *n.*

### 2.8 Calculation of circadian changes in functional networks

Lastly, we investigated whether there were circadian modulations in the functional connectivity networks. For this analysis, sleep and wakefulness were analyzed separately based on manual EEG sleep scoring. For both wakefulness and sleep, we calculated four metrics (the mean network strength and the three graph theoretical measures) as a function of time using ***Q*_300_** (see Section 2.5). Next, the values of each of these four metrics were associated with the 24-hour (circadian) clock time corresponding to the beginning of their ***Q*_300_** epoch. We did this for each subject in the dataset, thereby obtaining a distribution of values across subjects for each of the 1,440 circadian time-points (every minute on a 24-hour clock). We then calculated the mean across all subjects for each one-minute block of time; if there were less than five data points in the one-minute block, the data were considered insufficient and that time point was discarded. To assess statistical significance, we aggregated all data points from 11 AM to 1 PM as the “daytime” distribution and we aggregated all data points from 11 PM to 1 AM as the “nighttime” distribution. A Wilcoxon rank-sum test was used to determine whether the networks in the daytime and nighttime distributions exhibited significant differences.

## 3. Results

### 3.1 Subject demographics

In total, nineteen healthy subjects (15 female, 4 males) were recruited for the study between June 2017 and February 2019, and all were included in our analysis. The mean age of the subjects was 6.3 months (+/− 3.1 months, standard deviation). The recordings lasted 20.8 hours on average (+/− 7.8 hours, standard deviation).

### 3.2 Sleep is associated with stronger functional connections

Sleep was associated with stronger functional connections, as evidenced by the averaged connectivity networks ***Q**_w_* and ***Q**_s_* across all subjects (Figure 1A) and the individual subject results (Figure 1B). We also tested whether specific connection pairs were consistently stronger in wakefulness or sleep; to do this, we compared the distribution of connection strengths for one electrode pair during wakefulness to the distribution during sleep in a pairwise fashion (n=19 subjects). In 47 of the 171 possible connections, wakefulness revealed stronger connectivity values (Figure 1C, top) (two-tailed Wilcoxon sign-rank test, adjusted via Benjamini Hochberg procedure, adj. p<0.05). In 52 of the 171 possible connections, sleep connectivity values were significantly stronger than wakefulness (Figure 1C, bottom) (two-tailed Wilcoxon sign-rank test, adjusted via Benjamini Hochberg procedure, adj. p<0.05). The strongest connections in the averaged network were typically associated with sleep rather than wakefulness; although a number of connections were statistically stronger during wakefulness than sleep, these connections were typically weak, with strengths <0.04 (Figure 1D).

**Figure 1.**
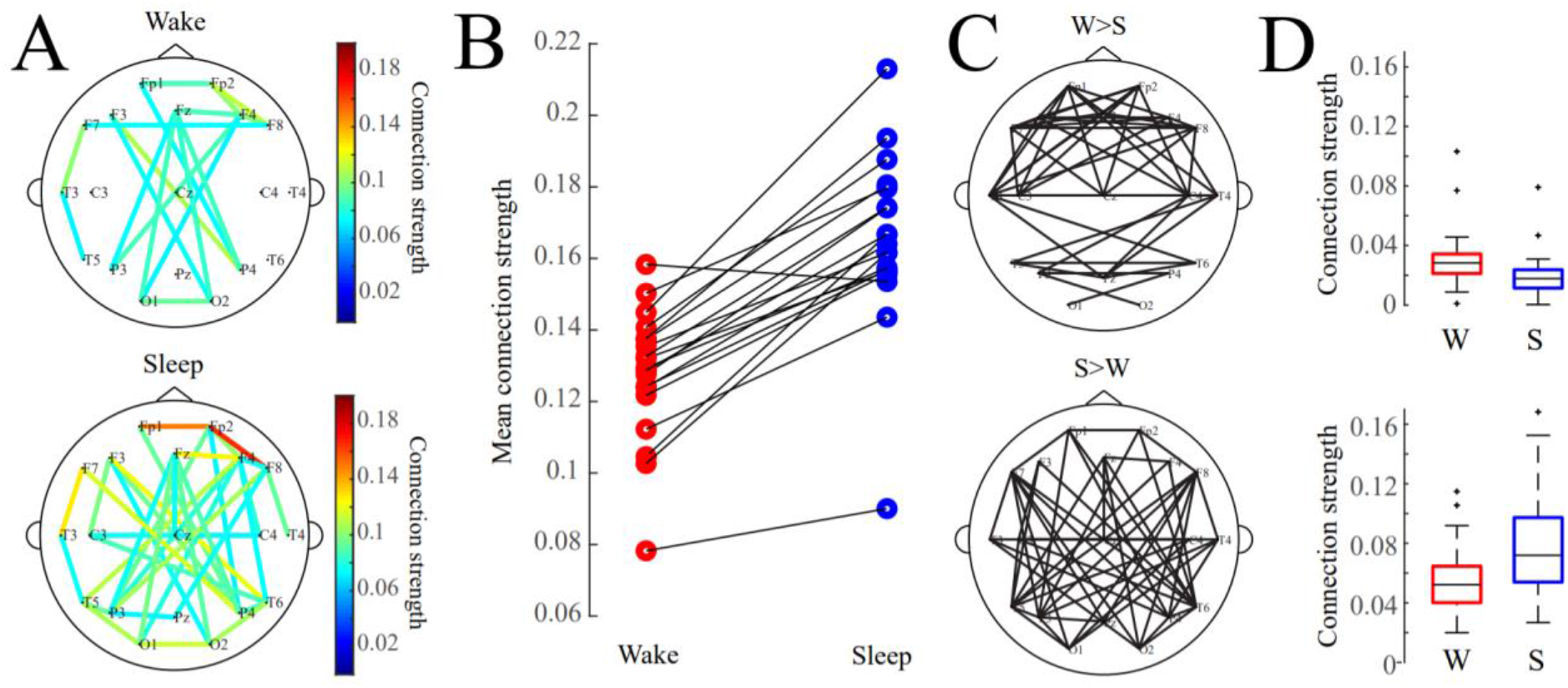
(A) Average functional connectivity networks for wakefulness and sleep. For visualization, an edge is drawn if the connection value exceeds an absolute threshold of 0.075. (B) Mean connection strength for individual subjects (calculated as the average strength of the strongest 10% of connections) is higher during sleep. (C) Network maps showing connections that were statistically significantly greater in wakefulness (top) or sleep (bottom). (D) Boxplots of mean connection strength for connections that were significantly different between wakefulness and sleep (shown in subfigure C).

### 3.3 Functional network structures are subject-specific

We then measured the similarity of the functional connectivity network structure within and across subjects, as well as within and across sleep/wake states (Figure 2). Across all subjects, the within-sleep distribution of correlation coefficients was statistically significantly higher than the within-wakefulness distribution, and both were significantly higher than the across-state distribution (Wilcoxon rank sum test, p<0.05). Additionally, within-subject across-state correlations (e.g. comparing subject 1 sleep to subject 1 wakefulness) were higher than across-subject within-state correlations (e.g., comparing sleep networks across all subjects). This indicates that, while state-specific functional network structure commonalities are seen across subjects, the network structures are also patient-specific. In other words, network structure is predicted to some degree by the sleep/wake state and to a greater degree by which subject it was recorded from.

**Figure 2.**
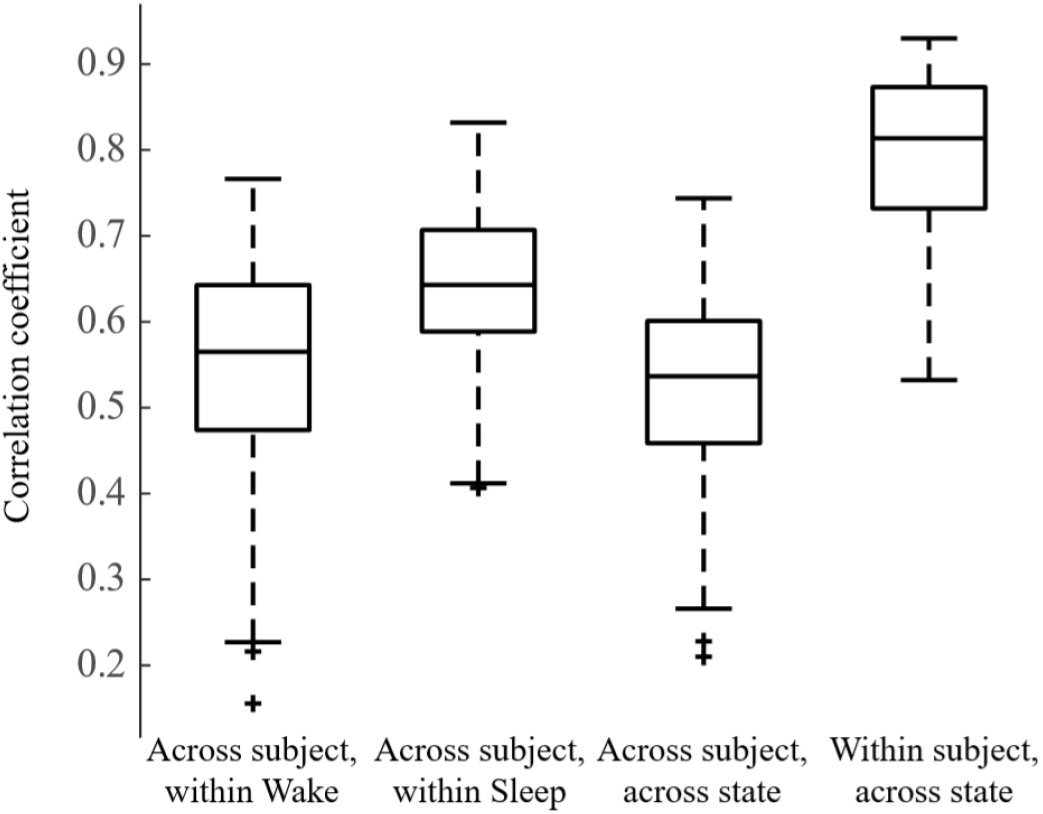
Box plot showing two-dimensional correlations within and across weighted connectivity matrices for each subject. We first compared networks across subjects within a state, e.g., compared Subject 1 Wake to Subject 2 Wake, and the analogous comparisons during sleep, n = 171 observations each. Then we compared across subjects and across states, e.g., Subject 1 Wake to Subject 2 Sleep, n = 171 observations. Lastly, we calculated the 2D correlation between the sleep and wake networks within single subjects, e.g., Subject 1 Wake to Subject 1 Sleep, n = 19 observations. All distributions are statistically significantly different from one another (Wilcoxon rank sum test, p<0.05).

### 3.4 Network structure is more segregated during wakefulness

We calculated three standard weighted graph theoretical measures on the average functional connectivity maps for sleep and wakefulness for each subject. Consistent with our finding that networks tended to be stronger during sleep than during wakefulness, the node degree was significantly greater in sleep than in wakefulness (Figure 3A) (Wilcoxon rank-sum test, p<0.05). The clustering coefficient was significantly greater in wakefulness compared to sleep (Figure 3B) (Wilcoxon rank-sum test, p<0.05). Lastly, we found the shortest average path length was not significantly different between sleep and wakefulness (Figure 3C) (Wilcoxon rank-sum test, p<0.05).

**Figure 3.**
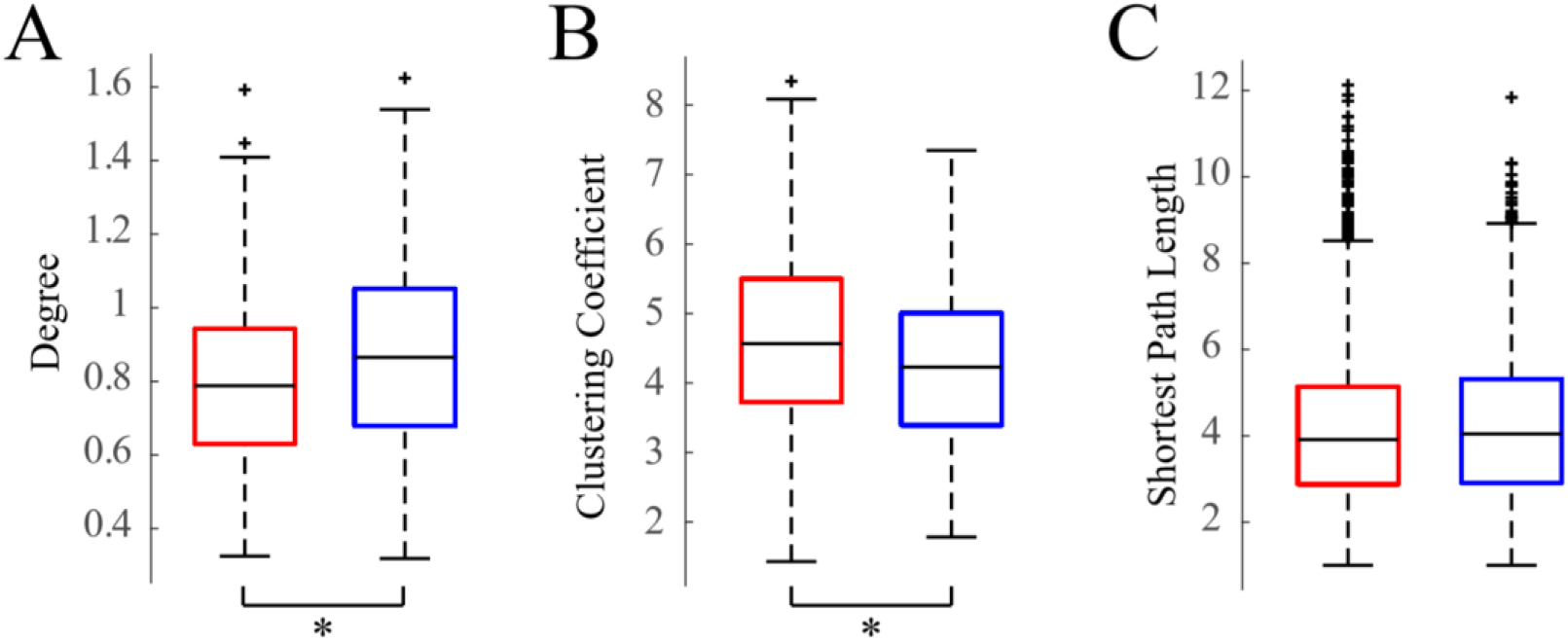
Boxplots of weighted graph theoretical measures. (A) Degree, (B) clustering coefficients, and (C) shortest path lengths of wakefulness (red) and sleep (blue) networks for all 19 subjects.

### 3.5 Functional networks are more stable in sleep than wakefulness

We then calculated the stability of each subjects’ functional connectivity networks within a given brain state (Figure 4). Here, stability was assessed using the correlation coefficient between networks calculated from independent windows of data. Higher correlation coefficients indicated greater similarity between networks and thus, higher stability. For each subject, we calculated the mean correlation coefficient for each window size and then calculated the 95% confidence interval for the mean correlation coefficients across all subjects. We found that sleep networks were significantly more stable than networks derived from EEG during wakefulness (Figure 4). The confidence intervals for the mean of the stability distributions did not overlap for any window size, indicating statistical significance over all tested window sizes.

**Figure 4.**
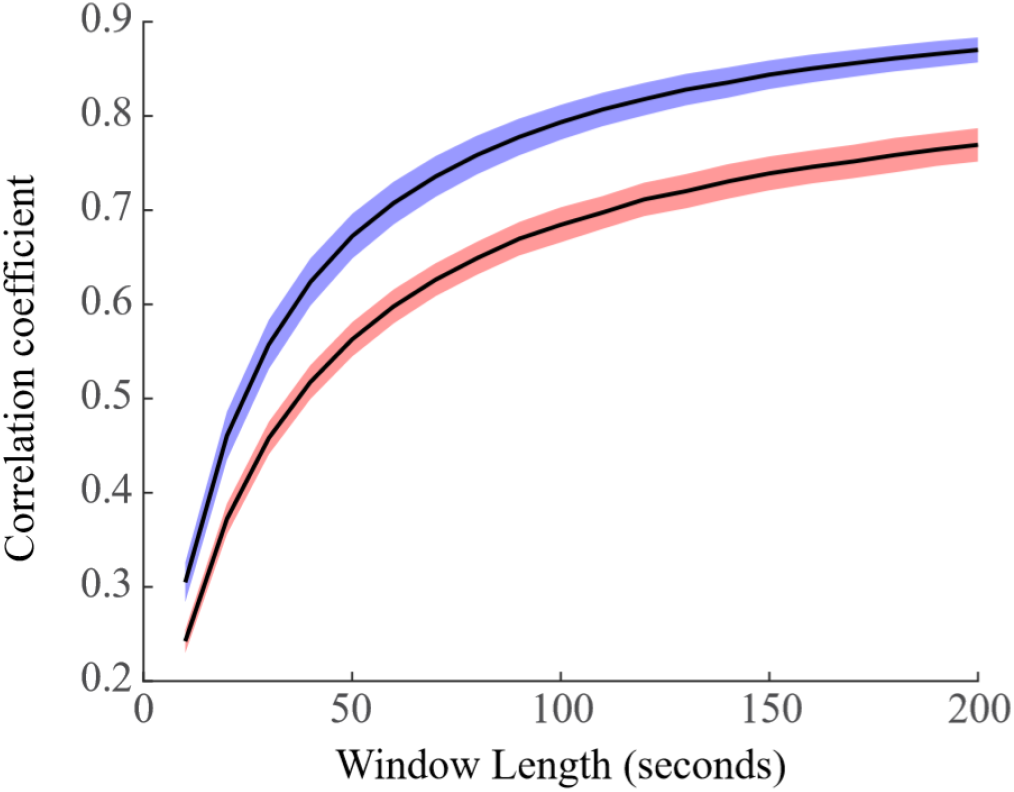
Stability of functional connectivity networks in wakefulness (red) and sleep (blue). We calculated the 2D correlation between independent averaged connectivity networks from windows of data of varying size. We found that sleep exhibited more stable networks, with nonoverlapping confidence intervals for the means for all window sizes.

### 3.6 Functional networks associated with sleep and wakefulness recur over hours and days

Our analysis showed that an individual subject’s functional network remains stable throughout each period of wakefulness or sleep. However, this analysis did not test whether the networks recur; that is, whether the functional network of one sleep period matches that of another sleep period within the same person. To address this, we used principal components analysis (PCA) to determine the latent variable that described the most variance in the connectivity data; we hypothesized that this variance would be due to transitions between wakefulness and sleep. If the networks remain stable for each state over multiple sleep/wake cycles, the time series of the first principal component, which signifies the weight of that component in the functional connectivity time series, should oscillate between two values.

We performed PCA on all functional connectivity network time series as outlined in Section 2.6. An example of a time series from the first principal component (PC1) is shown in Figure 5A, demonstrating the bimodal nature of the signal. This was also reflected in the histogram of the first principal component, which we fit with a two-component Gaussian mixture model (Figure 5B).

**Figure 5.**
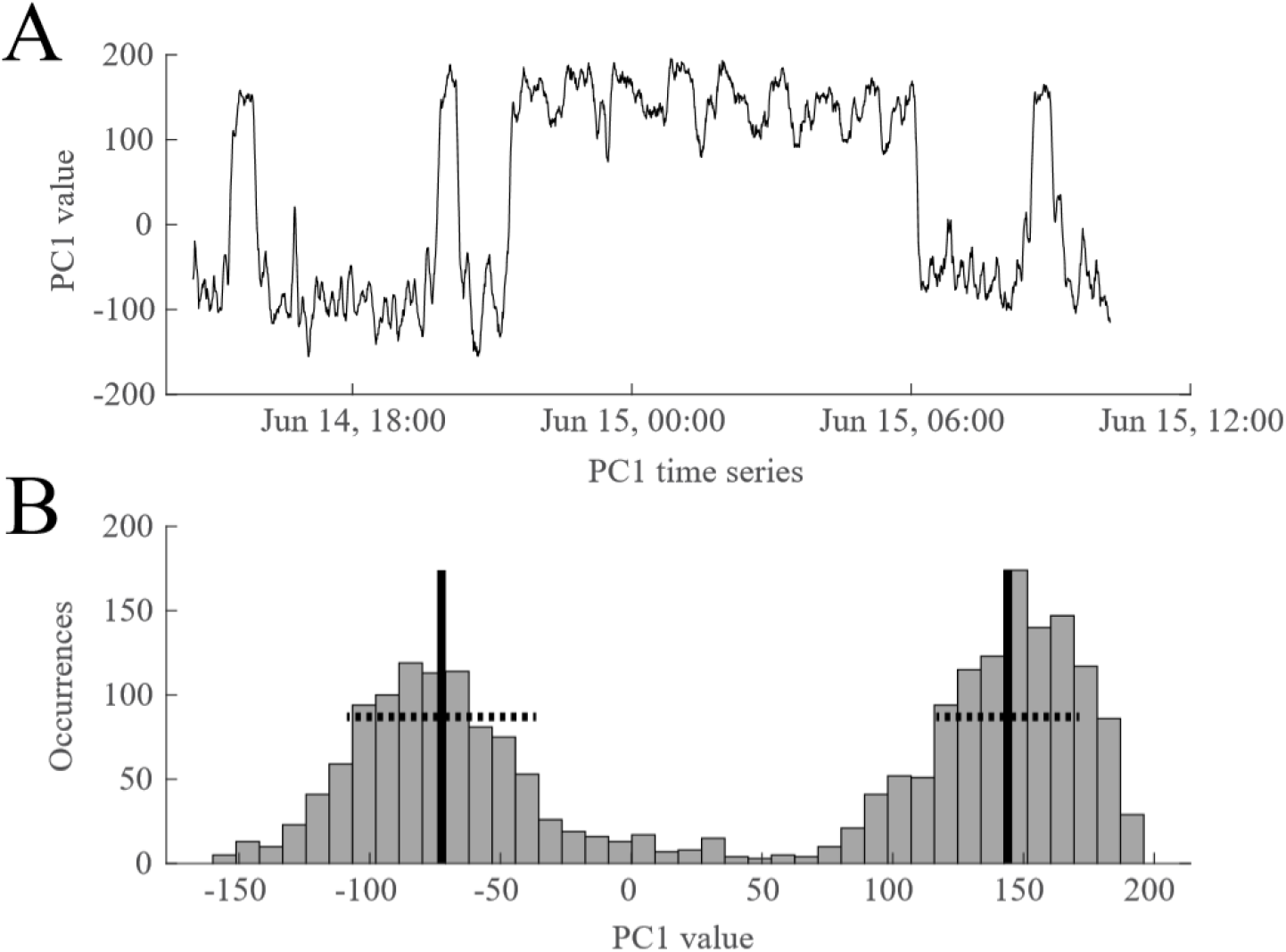
A representative example of the time course of the first principal component (PC1), reflecting how much weight is assigned to PC1 in the functional connectivity time series. (A) PC1 oscillates between two states during ~18 hours of EEG data. (B) The bimodal nature of PC1 is reflected in its histogram. A two-component Gaussian mixture model was derived from these values and used to classify the two states. The black vertical lines indicate the means of the two distributions and the dashed horizontal lines denote one standard deviation. Data are from Subject 1.

To determine whether the two reoccurring states evident in the PCA results corresponded to wakefulness and sleep, we compared the Gaussian mixture model output to visual sleep staging (Figure 6). Confusion matrices for each subject showed a high correspondence between the two states uncovered via PCA (defined by a threshold applied to the Gaussian mixture model) and the visually identified sleep and wake stages. Across subjects, the mean percentage of data consistent with this model was 80.2% and the median percentage was 91.2% (Table 1).

**Figure 6.**
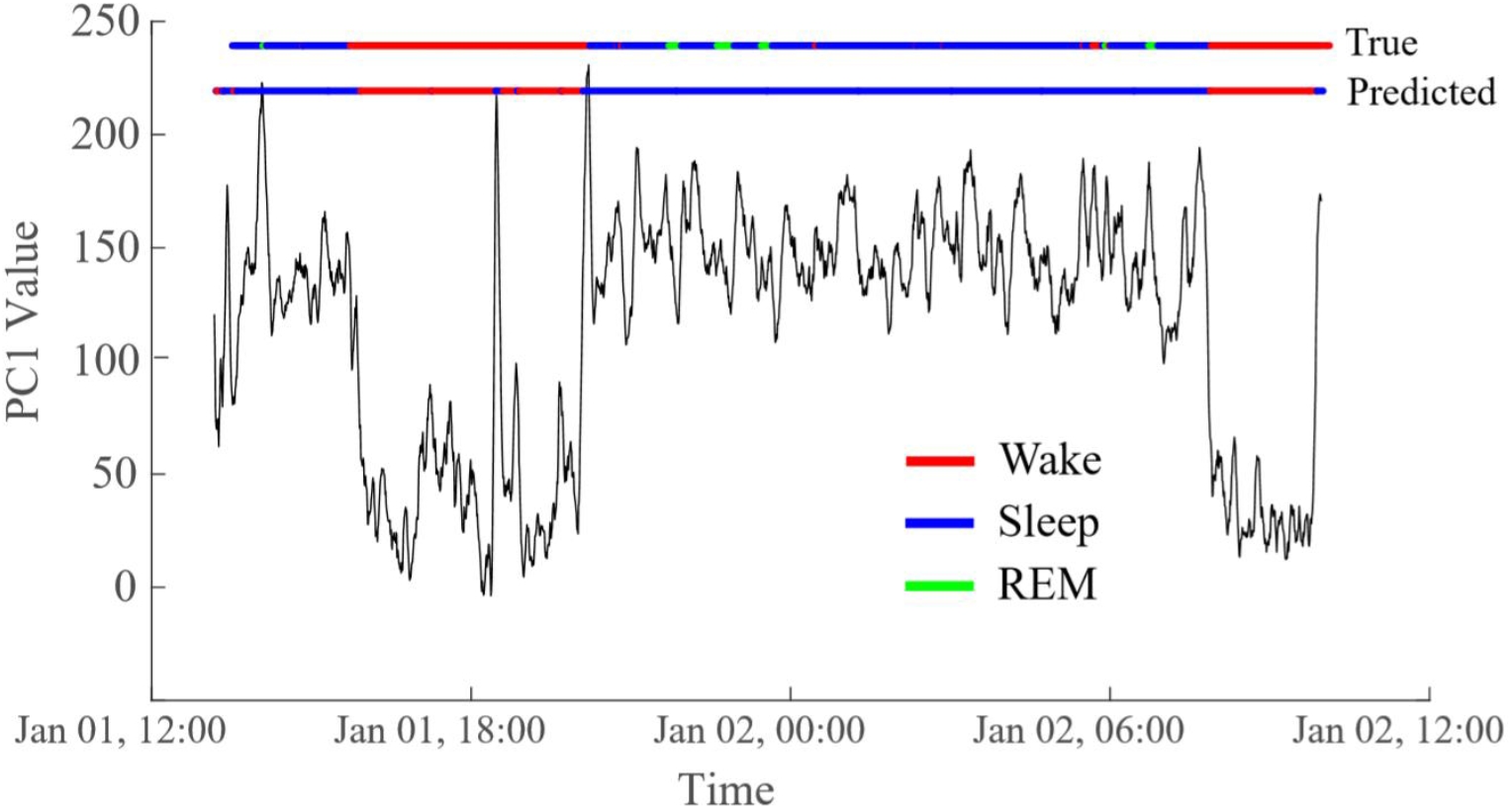
Automatic classification of two states using the PC1 time series matches visually-classified sleep stages. In this representative example, the accuracy is 95.1%. The top horizontal line (“True”) is colored to indicate the sleep stage based on visual markings. Red indicates the subject is awake, blue is non-REM sleep, and green is REM sleep. The bottom horizontal line (“Predicted”) reflects the automatic classification of sleep states based on a threshold applied to the Gaussian mixture model, with red representing wakefulness and blue representing sleep. For sensitivity, specificity, and accuracy calculations, REM was classified as sleep. Data are from Subject 5.

**Table 1.** Automatic identification of sleep/wake state for 19 subjects

|  | Scored Wake/<br>Classified<br>Wake (%) | Scored Sleep/<br>Classified<br>Sleep (%) | Scored Sleep/<br>Classified<br>Wake (%) | Scored Wake/<br>Classified<br>Sleep (%) | Percent<br>Correctly<br>Classified |
| --- | --- | --- | --- | --- | --- |
| **Subject 1** | 29.38 | 54.37 | 5.14 | 11.09 | **83.75** |
| **Subject 2** | 45.90 | 52.92 | 0 | 1.16 | **98.83** |
| **Subject 3** | 32.68 | 62.48 | 0 | 4.82 | **95.17** |
| **Subject 4** | 24.60 | 24.54 | 50.84 | 0 | **49.15** |
| **Subject 5** | 28.21 | 67.46 | 0.29 | 4.02 | **95.67** |
| **Subject 6** | 21.93 | 74.98 | 0.13 | 2.94 | **96.92** |
| **Subject 7** | 24.44 | 56.47 | 0.06 | 19.02 | **80.91** |
| **Subject 8** | 31.47 | 18.94 | 49.58 | 0 | **50.41** |
| **Subject 9** | 38.68 | 53.21 | 0.79 | 7.30 | **91.89** |
| **Subject 10** | 31.96 | 55.26 | 0 | 12.77 | **87.22** |
| **Subject 11** | 34.58 | 59.58 | 0.02 | 5.79 | **94.17** |
| **Subject 12** | 25.86 | 32.66 | 37.75 | 3.71 | **58.52** |
| **Subject 13** | 28.59 | 23.57 | 46.27 | 1.55 | **52.17** |
| **Subject 14** | 30.69 | 60.54 | 8.15 | 0.60 | **91.23** |
| **Subject 15** | 27.92 | 63.73 | 0 | 8.33 | **91.66** |
| **Subject 16** | 43.82 | 53.07 | 0.20 | 2.90 | **96.89** |
| **Subject 17** | 36.57 | 56.87 | 0.16 | 6.38 | **93.44** |
| **Subject 18** | 24.20 | 12.75 | 62.60 | 0.43 | **36.96** |
| **Subject 19** | 22.72 | 56.70 | 13.46 | 7.11 | **79.42** |

Subjects in which the percentage of correctly classified epochs exceeded 80% are highlighted in green (n=13 subjects). Subjects in which greater than 15% of the data were incorrectly classified are highlighted in red (n=6 subjects).

Thirteen of the 19 subjects exhibited persistent network patterns in wakefulness and sleep such that PCA captured the states over 80% of the time (Table 1). In subjects with lower classification accuracy based on PCA, it was common for the model to misidentify sleeping segments as wakefulness. Because our PCA analysis relied on the presence of both sleep and wakefulness to identify the two GMM distributions, the misidentification of sleeping segments often occurred in datasets in which the subject slept most of the recording and the algorithm misclassified the deeper sleep stages (N2 and N3) as wakefulness. On the other hand, in Subject 7 the model misidentified 19% of the data as sleep data when the subject was awake.

### 3.7 Long-term EEG functional networks exhibit circadian variation

When subjects were awake, the mean network strength was significantly decreased during the daytime and was increased at nighttime (Figure 7A, Wilcoxon rank sum test, p<1e-10). Similar patterns were seen for the network degree (Figure 7B) and the clustering coefficient (Figure 7C). The shortest path length showed the opposite trend (Figure 7D). Similar patterns were observed using data collected when subjects were sleeping, although the modulation over 24 hours was less dramatic, and the sleep measurements during the daytime exhibited high amounts of variability. In general, there was higher variance in the measurements when subjects were asleep during the daytime or awake at nighttime (in opposition to the expected circadian sleep patterns). However, all trends in graph theoretical measures were significantly different between daytime and nighttime hours (Figure 7B-D, Wilcoxon rank sum test, p<1e-10). This indicated that, in addition to the significant differences between sleep and wake functional networks, the time of day modulated the network within each state.

**Figure 7.**
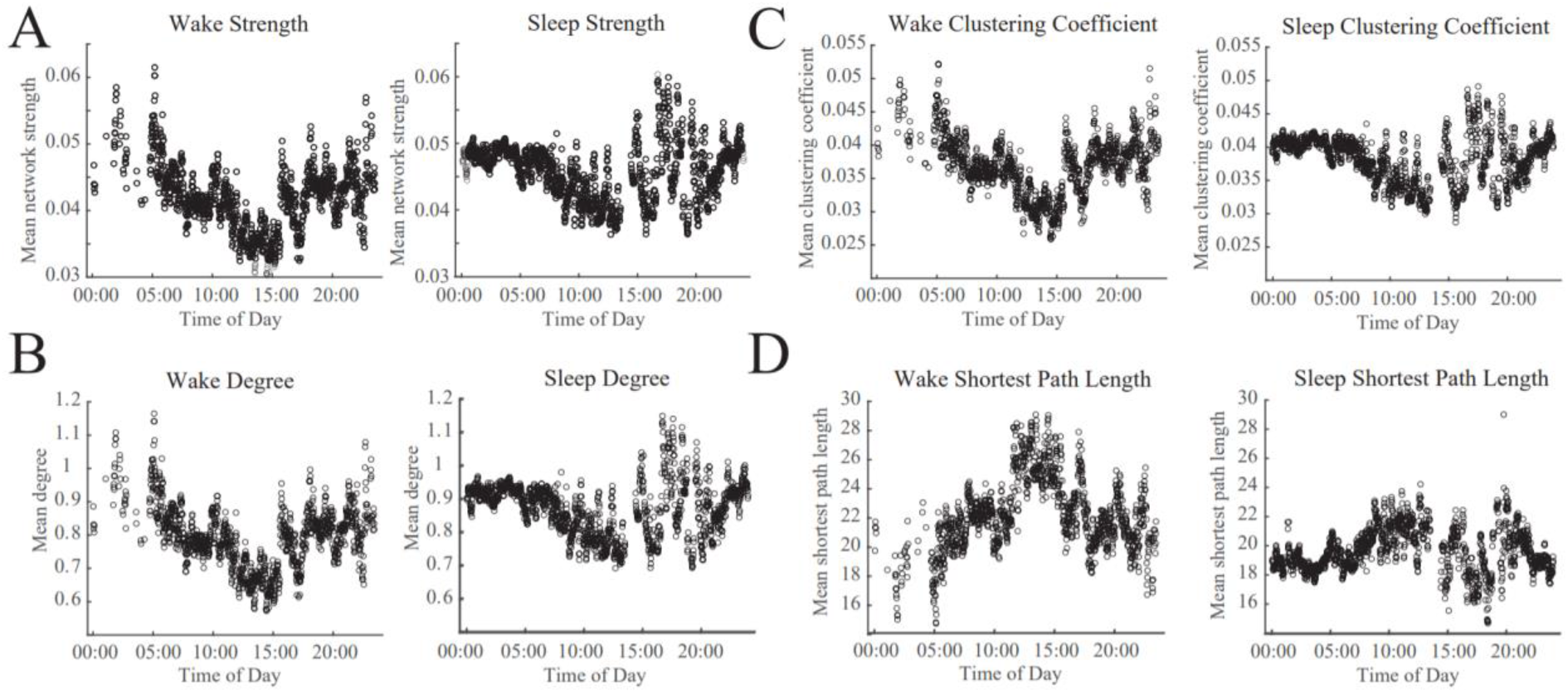
Circadian patterns emerge in both wakefulness and sleep for network strength and graph theoretical metrics. Twenty-four hour periodicities are shown for (A) the network strength, defined as the mean of all connections, (B) network degree, (C) clustering coefficient, and (D) the shortest path length. Each subfigure shows data recorded during wakefulness (left) and sleep (right).

## 4. Discussion

In this study, we report characteristics of functional connectivity networks based on long-term EEG recordings from 19 healthy infants. We first showed that functional connectivity networks associated with sleep and wakefulness exhibited statistically significant differences both in strength and structure. We also showed that, although transitions between sleep and wakefulness are associated with consistent changes to the network, the networks were highly individualized. Within-subject across-state network comparisons (comparing sleep to wakefulness) were more similar than comparisons across subjects within the same state. Further, state-specific networks recurred over multiple periods of sleep and wakefulness within each subject, and PCA enabled classification of the networks into these two states. Lastly, we showed that circadian rhythms significantly modulated network properties in a relatively stereotyped fashion. This suggests that the time of day during which a recording is obtained may significantly impacts measurements of functional connectivity, which bears relevance for both cognitive and clinical studies involving functional networks.

The analysis of infant EEG, which is visually distinct from adult EEG, is a unique aspect of this study. The vast majority of EEG functional connectivity studies focus on healthy adult data (Stevens, 2009); thus, this study fills a critical need by providing basic characteristics of healthy EEG-based functional networks as a baseline for studying conditions specific to the neonatal/infant period, including early-onset epilepsies and neuro-developmental conditions (Righi, Tierney, Tager-Flusberg, & Nelson, 2014; Shrey et al., 2018). Infant EEG poses unique advantages and disadvantages in comparison with adult recordings. On one hand, it enables study of circadian dynamics and network characteristics separately for sleep and wakefulness because infant sleep cycles do not always coincide with diurnal rhythms. On the other hand, the EEG patterns associated with sleep and wakefulness in infant EEG rapidly evolve as the infant grows and develops. Wakeful background activity in infant EEG is slower than adult EEG, and rhythms become faster with age (Fisch, 1999; Rowan & Tolunsky, 2003). The emergence of critical patterns, such as the posterior dominant rhythm and the mu rhythm, occurs at around 3 and 4-6 months of age, respectively (Laoprasert, 2011; Stern, 2005). Moreover, the structural connectivity in the infant brain is constantly changing and developing (Barkovich et al., 2006; Tymofiyeva et al., 2012), whereas structural connectivity in adults is relatively static. This age-dependence could partially explain the subject-specific nature of the networks observed in our study. Although researchers have advanced the study of the relationship between structure and function in the brain (Pernice, Staude, Cardanobile, & Rotter, 2011; Ponten, Daffertshofer, Hillebrand, & Stam, 2010), further work is needed to examine this relationship in the developing brain. Overall, we expect that the functional connections underlying infant neural activity will differ from adults.

We found that functional networks were stronger during sleep than wakefulness, and that they were less clustered when the subject was awake (Burroughs et al., 2014; Kuhnert et al., 2010; Mitsis et al., 2017). The networks were significantly stronger during sleep when compared to wakefulness in all but one subject (Figure 1B), however, we note that the effects of thresholding networks, even with a proportional threshold, is an active area of research and will require further investigation (Chapeton et al., 2017; Garrison et al., 2015). To reduce the bias introduced by thresholding the network graphs, we calculated graph theoretic properties of the networks on the weighted adjacency matrices. The degree was significantly higher in the functional networks derived from sleep, consistent with our finding of overall stronger networks in sleep. The clustering coefficient of the normalized networks was higher in awake networks, indicating greater functional segregation during wakefulness (Rubinov & Sporns, 2010; Watts & Strogatz, 1998). Interestingly, the shortest path length, calculated as the inverse of the normalized connection strength, was not significantly different between the two states, indicating similar levels of functional integration in wakefulness and sleep (Watts & Strogatz, 1998).

We found that the awake and sleep networks were more similar within a single subject than the awake or asleep networks across subjects. The subject-specific nature of these functional networks was also described in a long-term intracranial EEG study in adults (M. A. Kramer et al., 2011), as well as in a previous study by our group in a cohort of pediatric epilepsy patients (Shrey et al., 2018). This may indicate a need for a paradigm shift in the analysis of functional connectivity networks. Most functional network studies have focused on finding common networks and pathways that facilitate specific functions or the resting-state. However, the uniqueness of functional networks has become a recent area of investigation in the fMRI community and may deserve further attention in EEG functional network analysis (Chapeton et al., 2017; Chen et al., 2008; D’Esposito, 2019; Dubois & Adolphs, 2016; Fingelkurts & Fingelkurts, 2011; Finn et al., 2020; Gonzalez-Castillo et al., 2015; M. A. Kramer et al., 2011). A comparison of functional connectivity networks may require attention to both elements: the common pathways underlying the activity of interest, as well as the individuality of the subject’s unique functional network.

The stability of functional connectivity networks in long-term EEG is a function of the time scale used to measure them. We found that the binary connectivity matrices in one second epochs were highly variable, but stable networks were identified over the course of 200-500 seconds (Chu et al., 2012; Shrey et al., 2018). However, these networks become unstable again at longer time scales due to brain state transitions and circadian rhythms. This multi-level stability is assumed in our study, but further investigation is needed to define characteristic time scales of stability in functional connectivity networks in the human brain (Kuhnert et al., 2010). This is perhaps related to the concept that EEG amplitude modulations do not have a characteristic scale and exhibit a fractal nature (Hardstone et al., 2012; Linkenkaer-Hansen, Nikouline, Palva, & Ilmoniemi, 2001; R.J. Smith et al., 2017). This fractal nature may be transferred to functional networks (Lehnertz, Geier, Rings, & Stahn, 2017), mathematically suggesting that brain activity is changing in an organized way that may not have a characteristic time scale.

Several limitations of our study should be addressed in future investigations of healthy functional connectivity networks. First, our EEG studies were an average of 20.8 hours, and a limited number of recordings were longer than 24 hours. Thus, circadian patterns were assessed on a group level rather than an individual level. Future studies could include longer, multi-day EEG recordings to analyze true subject-specific assessments of circadian patterns. Second, we used an automatic algorithm to remove artifacts in our data, as it was infeasible to visually confirm all artifacts due to the long recording durations. Therefore, some artifacts may have escaped detection/removal while other artifact-free data may have been erroneously removed. This could have contributed to the differences seen in the wake and sleep networks because artifacts are more frequent during wakefulness. Lastly, although there are many advantages to analyzing data from a cohort of infants, the limited age range reduces the generalizability to other pediatric populations, and we did not have enough subjects to discern network properties specific to each age or developmental stage. Future studies should increase the number of subjects and broaden the age range.

The importance of this study lies in generating functional connectivity networks derived from long-term, normal EEG data. In addition to imparting knowledge of how physiological functional networks modulate throughout the day and within waking and sleep states, this will facilitate understanding of changes in network topology due to pediatric diseases such as epilepsy and autism (Righi et al., 2014; Shrey et al., 2018). Seizure forecasting in epilepsy has largely relied on prediction of seizure onset with several minutes of data, but it has been shown that modulations in functional networks due to physiological processes such as waking and sleeping can mask “pre-seizure” changes (Kuhnert et al., 2010; Mitsis et al., 2017; Schelter et al., 2011). Accounting for these physiological fluctuations in seizure prediction models may improve their accuracy and ultimately improve care for patients suffering from epilepsy.

## Acknowledgements

The authors thank the EEG technologists at the Children’s Hospital of Orange County (CHOC) for their help in acquiring the EEG data. The authors also thank Dr. Michael Nunez and Derek Hu for helpful discussions regarding the manuscript. This study was funded by a CHOC PSF Tithe grant.

